# Charge influences substrate recognition and self-assembly of hydrophobic FG sequences

**DOI:** 10.1101/090159

**Authors:** Wesley G. Chen, Jacob Witten, Scott C. Grindy, Niels Holten-Andersen, Katharina Ribbeck

## Abstract

The nuclear pore complex controls the passage of molecules via hydrophobic phenylalanine-glycine (FG) domains on nucleoporins. Such FG-domains consist of repeating units of FxFG, FG, or GLFG sequences, which can be interspersed with highly charged amino acid sequences. Despite the high density of charge exhibited in certain FG-domains, if and how charge influences FG-domain self-assembly and selective binding of nuclear transport receptors is largely unexplored. Studying how individual charged amino acids contribute to nuclear pore selectivity is challenging with modern *in vivo* and *in vitro* techniques due to the complexity of nucleoporin sequences. Here, we present a rationally designed approach to deconstruct essential components of nucleoporins down to 14 amino acid sequences. With these nucleoporin-based peptides, we systematically dissect how charge type and placement of charge influences self-assembly and selective binding of FG-containing gels. Specifically, we find that charge type determines which hydrophobic substrates FG sequences recognize while spatial localization of charge tunes hydrophobic self-assembly and receptor selectivity of FG sequences.

## Introduction

The nuclear pore complex (NPC) is a megadalton structure that controls the exchange of material between the nucleus and cytoplasm (1–3) through a combination of passive and facilitated diffusion. Above a ~30 kDa cutoff, proteins require complexation with nuclear transport receptors (NTRs) to efficiently translocate at rates of nearly one thousand molecules per second (4–7). This fast translocation rate relies on transient interactions between hydrophobic phenylalanine-glycine (FG) domains on intrinsically disordered FG-containing nucleoporins (FG-nups) and hydrophobic patches on NTRs (1,5,8–20). Without such assistance of NTRs, proteins remain excluded from the NPC. Despite the necessity of FG-domains (which contain repeating units of FG sequences such as FxFG or GLFG) for facilitated diffusion and self-assembly of the selective matrix, how hydrophobic FG-domains exhibit such selectivity to specific hydrophobic domains on NTRs and what parameters can tune such molecular recognition necessary for facilitated transport remain open questions. Sequence analysis and molecular dynamic simulations predict that the biochemistry and charge surrounding individual FG sequences should play an essential role in FG-mediated molecular recognition and therefore determine the organization and selectivity of the nuclear pore (4,6,21–24). However, due to the complexity and redundancy of FG-nups within NPCs, a systematic biochemical dissection of the amino acid space surrounding individual FG sequences has remained an experimental challenge.

In this study, we investigate if and how electrostatic interactions surrounding FG sequences tailor both the self-assembly of FG sequences and the selective recognition of hydrophobic substrates. For systematic dissection of how charge type and localization influence FG-mediated selectivity, full FG-domains are too degenerate and complex to provide insight into how single amino acids contribute to selectivity. To overcome this challenge, we established a novel approach that uses short peptides. The use of peptides and polypeptides has previously been essential in design of other minimalistic polymer-based biomaterials such as extracellular matrices (25), silk proteins (26,27), and elastin-like polypeptides (28–30). While peptides may not recapitulate all properties of the original protein, they allow for mechanistic dissection of structure and function at the single amino acid level. Here, peptides can be designed to represent the essential components of charged FG-nups, allow for single amino acid substitutions, and can be engineered to self-assemble into hydrogels mimicking the high density of FG-domains found in certain FG-nups. With this peptide system, we show how the amino acids surrounding FG sequences has a significant impact on hydrophobic interactions necessary for molecular recognition of substrates and self-assembly. Choice of anionic and cationic residues is able to reverse the selective recognition of the FG-based gels. This allows for FG sequences to discriminate between substrates exhibiting various types of hydrophobic domains using a combination of hydrophobic and electrostatic interactions. Moreover, spatial localization of charge determines the degree to which electrostatic interactions influence hydrophobic interactions with FG sequences. Lastly, we find that the choice of charged residues such as lysine, arginine, glutamic acid, and aspartic acid have different contributions to the self-assembly of FG sequences while electrostatic properties dominate for molecular recognition.

## Results

As a model FG-nup to base our FG-domain mimicking peptides on, we chose the yeast nucleoporin Nsp1 as it is essential in *Saccharomyces cerevisieae* (31) and contains a repeating subsequence (284-553) with a high density of charge (Figure 1A). Sequence analysis of 15-repeats of FG-domains in Nsp1^284-553^ reveals a non-uniform distribution of charge in the sequence space separating FG sequences (here, specifically, we use the FSFG sequence). Cationic residues can be found near the center and edges of the repeat whereas anionic amino acids reside at least three positions away from FSFG sequences (Figure 1B). Moreover, we find several highly conserved lysines (K) situated two to three amino acids away from the FG-domain at positions 6 and 17 and a conserved glutamic acid (E) at position 9 (Figure 1A). The non-uniformity in conserved charge distributions and amino acid identity suggest that charge and biochemical properties of amino acid sidechains may play an essential role in governing how neighboring FG sequences respond to environmental substrates and that significant function can be encoded in the non self-assembling domains of FG-nups.

**Figure 1:**
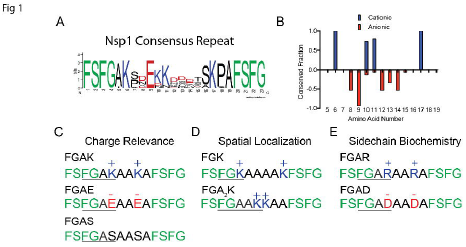
Identification of conserved repeat sequences and design of simplified FG self-assembling peptides A) Conserved sequence identification of the C-terminal end of essential yeast nucleoporin Nsp1. B) Conservation of charge between FSFG domains in Nsp1 sequences. Positive/negative values refer to the presence of cationic or anionic residues. Amino acid numbers correspond to position in the consensus repeats as seen in Figure 1A. C) Simplified peptides consisting of 14 amino acids were designed with the sequence structure FSFGAXAAXAFSFG where X represents the substituted amino acids lysine (K), glutamic acid (E), or serine (S) to determine how presence of charge affects FG-mediated selectivity. C) Designed peptides where the lysines are moved either immediately adjacent (FGK) or placed three away (FGA_2_K) from FSFG domains to test how spatial localization of charge influences FG selectivity. D) Class of peptides where lysines (K) are substituted with arginine (R) or aspartic acid (D) to determine how biochemistry affects FG-mediated selectivity and self-assembly.

For testing each of the parameters of charge type, placement, and sidechain biochemistry and how they affect FG function, we synthesized the consensus Nsp1 peptide sequence and also a simplified 14 amino acid version. The deconstructed peptides consist of two terminal FSFG sequences with neighboring charged or neutral amino acids. To establish how the presence of charge affects FG-mediated selectivity, we designed the peptide sequence FSFGAXAAXAFSFG, where X is either a lysine (K), glutamic acid (E) or a neutral serine (S) as seen in Figure 1C. Since lysine overall is the most highly conserved residue in this wildtype Nsp1 subsequence, we use the lysine-containing peptide as the reference for all other experimental comparisons (termed FGAK peptide). To determine if the selectivity properties of FG-domains could be reversed by switching neighboring charge types, we synthesized the anionic peptide FGAE. As a control of how the presence of charge influences FG function, we synthesized the neutral FGAS-peptide in which the lysines have been converted to serines. To test how the spatial localization of lysines affect FG function, we designed two variants of FGAK in which the lysines have been placed directly adjacent to the FG-domain or separated by two alanines (termed FGK and FGA_2_K, respectively, Figure 1D). Last, it appears that only certain charged amino acids such as lysines and glutamic acids (E) exhibit 100% conservation at positions 6, 9, and 17 (Figure 1A), suggesting amino acid biochemistries may also regulate FG-based molecular recognition. To test how chemical structures of the amino acid sidechains affects structure and function of FG-domains, we designed a third class of peptides in which the lysines are replaced with cationic arginines (R, FGAR) or aspartic acids (D, FGAD) as seen in Figure 1E. With these 14-amino acid nucleoporin-based peptides, we now analyze how single amino acid substitutions can alter the selective binding and self-assembling capabilities of individual FG-domains.

## Characterization of charge presence on FG-mediated self-assembly

Before studying the selective recognition of the FG-based peptides, we first establish if the peptides are capable of forming gels at a concentration relevant to intact nuclear pore complexes. By measuring the mechanical properties of the peptide solutions, we can further understand how charge affects FG-domain interactions. We used a concentration of 2% (w/v) for each of the peptides, which corresponds to 28 mM concentrations of FG sequences, a value that is well within estimations of FG sequences found in densely packed NPCs (18,32,33). To quantify gelation, we measure the stiffness of the resulting material using small amplitude oscillatory frequency sweeps. We report the storage (G’) and loss (G”) modulus of the peptide solution and specify that a gel forms when G’ is greater than G”, which indicates successful self-assembly of a stable network of peptides. The consensus peptide sequence of Nsp1 was unable to form a gel and flowed when inverted, so it was not suitable for further analysis. For the simplified peptides, we first establish that the reference peptide FGAK is able to form hydrogels with a stiffness of 10^4^ Pa across the frequencies tested (Figure 2A). To ensure that the FG sequences are responsible for the self-assembly process, we converted the hydrophobic phenylalanines to serines (SGAK). With this substitution, the peptide solution remains in an aqueous phase and no longer exhibits a dominant storage modulus (Supplementary Figure 1), suggesting that phenylalanine provides the necessary hydrophobic interactions for self-assembly. Reversing the charge to glutamic acids (FGAE peptides) reveals that the gel-forming properties are maintained with stiffness between 5-9 KPa at the frequencies tested (Figure 2B). To determine if the charge is responsible for maintaining the hydrated state of the gel in the self-assembly process, we compared the material to the solution of neutral FGAS peptides. Without the charge, FGAS precipitates out of solution (data not shown). Together, these results indicate that the presence of charge is essential in maintaining a hydrated network of FG sequences in self-assembly. However, too much charge increases the solubility and prevents gelation, as seen with the consensus sequence peptide. Without the presence of electrostatic repulsion within the gels, the network collapses and forms a precipitate. Moreover, for this class of designed peptides, FG sequences are essential for gelation, which is the same as for intact FG-nups (9,10).

**Figure 2:**
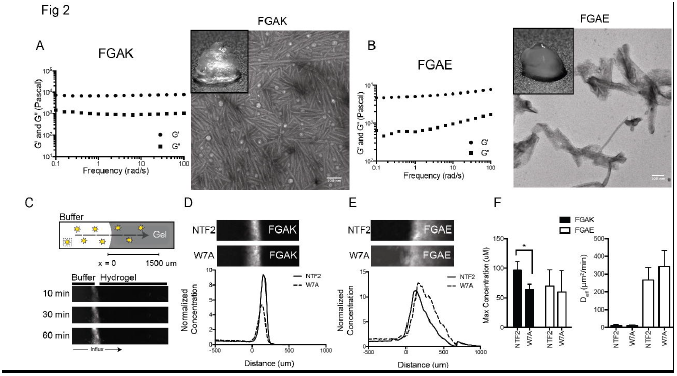
FGAK and FGAE self-assembly and selectivity of native transport receptor NTF2 A) Frequency sweep of FGAK gel with G’ (storage) and G” (loss) moduli reported at 2% (w/v) with TEM image of self-assembled peptides and macroscopic gel (inset). B) Frequency sweep of FGAE gel with G’ (storage) and G” (loss) moduli reported at 2% (w/v) with TEM image of self-assembled peptides and macroscopic gel (inset). C) Schematic of slab 1-D transport system within a sealed capillary providing no flux boundary conditions. Example diffusion of fluorescent molecules is depicted at 10 minute, 30 minute, and 60 minute timepoints. The interface of the gel is depicted by the increase in fluorescence signal between the buffer and hydrogel. D-E) Transport of representative fluorescently labeled NTF2 and W7A proteins into FGAK and FGAE gels, respectively. The fluorescent images and corresponding concentration profiles are at the 5-hour time point. F) The max concentration at the interface and effective diffusion coefficient are quantified for both NTF2 and W7A in FGAK and FGAE gels. All error bars are standard deviations of at least three independent replicates. * indicates p< 0.05 using unpaired Student’s t-test.

## Transport of NTF2 is asymmetric in cationic and anionic FG-based gels

To understand if the FG-based peptide gels are able to recapitulate the selective recognition properties of the native FG-nups, we test for their ability to enrich and select for intact nuclear transport receptors (NTRs) in an FG-dependent manner. As a model NTR, we choose NTF2 as it contains an essential tryptophan (W7) that is required for binding to the FG-domains. By substituting the tryptophan for an alanine (W7A) to ablate FG-binding capabilities(14), we can determine if the FG-domains are available in the gel for NTR binding. First, we determine if the positively charged FGAK gel is able to preferentially select for NTF2 compared to the W7A mutant. To prepare gels for selective transport assays, we dissolve FGAK peptides at 2% (w/v) as before and load the material into capillaries. Fluorescently labeled NTF2 or W7A is then injected into the capillary and sealed to create a 1-D diffusion chamber where the diffusion profile is monitored for up to 5 hours at one-minute intervals (Figure 2C). To quantify the selective binding of the various transport receptors into gels, we estimate the effective diffusion coefficient of the receptors in the gel using the time-evolving fluorescent profiles and the maximum accumulation of the reporters at the interface as metrics for determining how well the gel is able to differentiate between native NTF2 and W7A. Lower effective diffusivities and higher accumulation indicate stronger interactions with the gel.

As seen in Figure 2D and F, the FGAK gel accumulates NTF2 at higher concentrations at the gel interface compared to the W7A mutant. While accumulation at the interface is different, we find the effective diffusivities to be similar (Figure 2F), suggesting that the favorable hydrophobic interactions are weak and reversible, which is consistent with previous models of nuclear pore transport. These experiments show that FGAK has a selective binding preference for NTF2 over the W7A mutant, indicating that FG-domains are available and functional as a binding domain. It also appears that the short peptide gels cannot recapitulate the unique facilitated diffusion properties of the intact nuclear pore. For instance, native NTRs that are overall anionic and hydrophobic are predicted to bind to and transport into overall cationic nucleoporin gels (21,22). Since the peptides form fibers and sheets (Figure 2A-B) that are not observed in native nucleoporin assemblies (24), they likely do not form the requisite saturated disordered matrix that allows for exclusion of inert molecules that typically cannot transport through NPCs (9). Nevertheless, the designed FGAK peptide has the sensitivity to recognize single hydrophobic amino acid differences between NTF2 and W7A, which is a necessary first step for selective hydrophobic transport.

To test if binding to NTRs is sensitive to charged residues neighboring the FG domain, we determine if FGAE is able to recapitulate the same selective properties of FGAK, despite the reversal of charge. As seen in Figure 2E-F, FGAE selects for NTF2 and W7A equally at the interface and there is no significant difference in diffusion coefficients. These results indicate FGAE is unable to differentiate between the native and mutant form of NTF2. Since FGAK and FGAE exhibit different selective properties for NTF2 and W7A, we further investigate if the asymmetry results from a structural issue or a charge based selective recognition. Figures 2A-B show that FGAK and FGAE gels exhibit structural differences at the microscopic level (sheets vs. fiber assembly) and macroscopic level (inset). It is possible that due to structural differences, the FG sequences are not exposed and unavailable for binding by NTF2 in the FGAE gel. To test if hydrophobic domains are accessible, we use Nile Red dye, which fluoresces within hydrophobic environments. Supplementary Figure 2 shows that Nile Red fluorescence is detected in both FGAK and FGAE gels, indicating that the uncharged hydrophobic dye is able to interact with the FG-domains within the peptide-based gels despite the structural variation. These results suggest that the lysine and glutamic acid are important in regulating the selective properties of the FG sequences from a charge interaction perspective, as opposed to simply altering the gel’s microstructure.

## Presence of charge regulates the selective recognition of FG-domains

Although NTRs such as Importinβ and NTF2 have well-characterized specificity for particular FG-nup sequences, intact receptors contain multiple binding pockets with varying affinities and charge distributions, or require dimerization for function, which complicates systematic analysis of charge contribution. Hence, as a replacement for complex NTRs, we designed fluorescent peptide reporters with defined spatial arrangements of charged and non-polar amino acids to systematically test the contribution of charge in hydrophobic selectivity (Figure 3B). The first two reporter peptides contain three phenylalanines for a hydrophobic tail that enable binding to the FG sequences, but one contains an adjacent anionic sequence composed of glutamic acids (termed Hydrophobic (−)), while the other contains cationic lysines (Hydrophobic (+)). As controls, we engineered two more reporters where phenylalanines were converted to hydrophilic asparagines (N), and termed Hydrophilic (− or +) and should not interact with FG-domains. To confirm that the synthetic hydrophobic reporters are able to interact with the aromatic phenyl group in FG domains in the gels independently of their charged domains, we used phenyl-sepharose columns to test for hydrophobic interactions. This assay has previously been successful in isolating intact NTRs from cell lysate extracts (12). As seen in Supplementary Figure 3, both hydrophobic reporters show an increase in retention time compared to their hydrophilic counterpart, indicating that the hydrophobic interactions are occurring between the hydrophobic reporters and the phenyl groups on the sepharose independent of the displayed charge.

**Figure 3:**
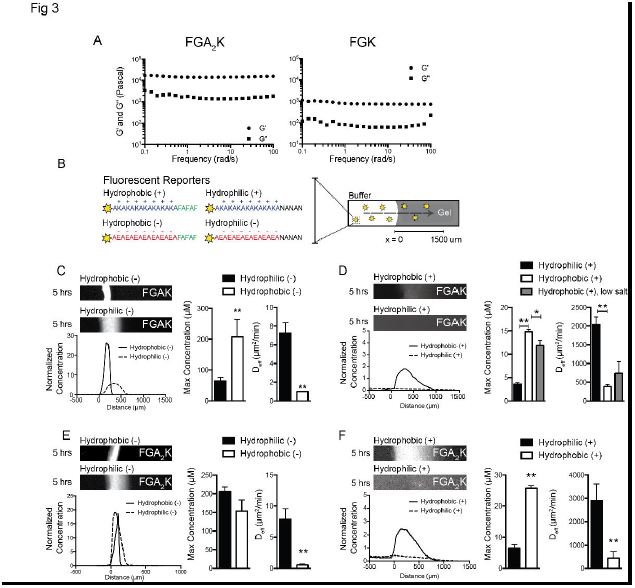
Effects of spatial localization of charge on FG-mediated self-assembly and selectivity A) Frequency sweep of FGA_2_K and FGK gels with G’ (storage) and G” (loss) moduli reported at 2% (w/v). B) Schematic of the four fluorescent reporters that contain a charged domain and a hydrophilic (N) or hydrophobic (F) tail. Fluorescent reporters are loaded into capillaries containing the gel of choice where diffusion is monitored for up to 5 hours. C) Representative fluorescence images of Hydrophobic (−) and Hydrophilic (−) reporters into FGAK gel with the corresponding concentration profile, max concentration at gel interface, and estimated effective diffusion coefficient. D) Representative fluorescence images of Hydrophobic (+) and Hydrophilic (+) reporters into FGAK gel with the corresponding concentration profile, max concentration at gel interface, and estimated effective diffusion coefficient. An additional low salt condition was added (20 mM NaCl) for Hydrophobic (+) reporters as a comparison to standardized (200 mM NaCl) conditions for max concentration and effective diffusion estimation. E) Representative fluorescence images of Hydrophobic (−) and Hydrophilic (−) reporters into FGA_2_K gel with the corresponding concentration profile, max concentration at gel interface, and estimated effective diffusion coefficient. F) Representative fluorescence images of Hydrophobic (+) and Hydrophilic (+) reporters into FGA_2_K gel with the corresponding concentration profile, max concentration at gel interface, and estimated effective diffusion coefficient. All error bars are standard deviations of at least three independent replicates. * indicates p<0.05. ** indicates p<0.01 between hydrophobic and hydrophilic reporters using unpaired Student’s t-test.

Using the FGAK gels, we first tested if the two hydrophobic fluorescent reporters demonstrate different selective uptake into the gel. FGAK was dissolved at 2% (w/v), loaded into glass capillaries, and bathed in a fluorescent reporter solution to test for selective binding. Reporters were loaded into capillaries at 10 μM concentrations and interactions were monitored for five hours at the buffer-gel interface. To quantify the selective binding of the various reporters into the respective gels, we again estimate the effective diffusion coefficient of the reporters in the gel and the maximum accumulation of the reporters at the interface as metrics for determining how well the gel is able to differentiate between hydrophobic and charged domains in the reporters.

Figure 3C shows that the cationic FGAK gel interacts with the Hydrophobic (−) reporters, with significant accumulation inside the gel. Quantifying the maximum concentration at the interface, we see that there is approximately a 2-fold increased enrichment compared to the Hydrophilic (−) reporter, which does not contain the hydrophobic tail. This result suggests that FG sequences in FGAK recognize reporter phenylalanines when the reporter exhibits negative charge. The contribution of hydrophobic interactions is further corroborated by the strong reduction in effective diffusion coefficient from 7 μm^2^/min for Hydrophilic (−) down to below 1 μm^2^/min for Hydrophobic (−), indicating that the hydrophobic interactions are able to induce tighter binding in the context of electrostatic attraction. In fact, the interactions are so strong that the Hydrophobic (−) reporters essentially do not diffuse over the five-hour period analyzed, suggesting that the binding is irreversible. In contrast, when the reporters contain a cationic tail, we see that Hydrophobic (+) and Hydrophilic (+) reporters diffuse into the gel with coefficients of 380 μm^2^/min and 2000 μm^2^/min, respectively. The Hydrophobic (+) reporters are able to bind 1.5 times above the original bath concentration at the interface whereas Hydrophilic (+) reporters equilibrate with no discernible partitioning (Figure 3D). These results show that hydrophobic interactions can occur under electrostatically repelling environments, but the interactions are much weaker compared to the Hydrophobic (−) reporters. Moreover, by lowering the salt concentrations from 200 mM NaCl to 20 mM to decrease electrostatic screening and increase electrostatic repulsion, the accumulation at the interface decreases and the diffusion coefficient shows a trend toward an increase (Figure 3D), indicating that the repulsion weakens the overall strength of hydrophobic interactions with FG-domains. These results show that the lysines confer upon FGAK gels the striking ability to distinguish between two substrates that contain the same hydrophobic domain but with different surrounding charge types. In particular, the lysines allows for increased binding to hydrophobic domains with neighboring anionic residues.

The analysis of the Nsp1 repeat consensus sequence (Figure 1A-B) suggests that the conserved location of lysines within 2-3 amino acids from the FG sequences may be relevant for FG-mediated molecular recognition. To determine how the spatial localization of lysines affects FG sequence differentiation between various hydrophobic domains, we tested if placing lysines immediately adjacent to FG sequences (FGK), or moving them 3 amino acids away (FGA_2_K), would affect the sequence’s selectivity compared to the original arrangement in the FGAK peptide. Using the same conditions as before, both peptides were dissolved at 2% (w/v). While FGA_2_K peptides readily formed a stiff material and could be tested for selective uptake, the FGK peptide gel is two orders of magnitude less stiff, flowed when inverted, and dispersed when loading fluorescent reporters (Figure 3A). Therefore FGK could not be analyzed for transport and the rest of the analysis will be a comparison on diffusion and accumulation between only FGAK and FGA_2_K. As previously shown in Figure 3C and D, the FGAK gel selectively enriched for the Hydrophobic (−) reporter but failed to uptake the Hydrophobic (+) reporter to the same degree. When comparing the two anionic Hydrophobic (−) and Hydrophilic (−) reporters in FGA_2_K, the diffusivity of Hydrophobic (−) is an order of magnitude lower compared to Hydrophilic (−), showing that the combination of hydrophobic and electrostatic attraction synergizes for strong binding. However, the max accumulation at the interface for the two anionic reporters is similar, indicating FGA_2_K gel is unable to differentiate between the two reporters to the same degree as the original FGAK gel (Figure 3E). For cationic reporters, the Hydrophobic (+) peptide accumulates within the gel and has an order of magnitude lower diffusivity compared to the Hydrophilic (+) reporter. Moreover, the accumulation of Hydrophobic (+) reporters for FGA_2_K gels is higher compared to FGAK gels (Figure 3F). These data show that placing lysines within 2 amino acids of FG-domains (FGAK), approximately corresponding to the 1 nm Debye length at physiologically relevant salt concentrations, such as in our experimental conditions, enables hydrophobic selectivity of substrates with opposite charge. Placing lysines 3 amino acids away (FGA_2_K) reduces the contribution of electrostatic interactions in determining hydrophobic FG-mediated selectivity. As a result, by increasing the distance between charged residues and FG sequences, the FG sequences becomes less dependent on the electrostatic profile surrounding the hydrophobic substrates during selective transport.

## Charge reversal flips selectivity of FG-domains

Since lysines in the FGAK gel help FG sequences differentiate between reporters containing anionic or cationic hydrophobic domains, we next asked if FG-mediated recognition could be reversed by replacing lysines with anionic glutamic acids (FGAE). Using the same class of fluorescent reporters, we find that the selectivity of FGAE gels is reversed from FGAK-gels (Figure 4A-D). Again, the selectivity is a function of both hydrophobic and electrostatic interactions as the Hydrophilic (+) reporters accumulated 2-fold less at the interface compared to Hydrophobic (+) reporters (Figure 4D). Moreover, the diffusion coefficient for Hydrophobic (+) is an order of magnitude lower than for Hydrophilic (+). Conversely, both Hydrophobic (−) and Hydrophilic (−) reporters did not interact significantly with the gel and exhibited similar effective diffusion coefficients, showing that electrostatic repulsion prevents hydrophobic interactions with FG sequences. Together, these data suggest that charge proximal to FG sequences can tune hydrophobic selectivity and that charge is essential in determining which hydrophobic moieties FG sequences can recognize.

**Figure 4:**
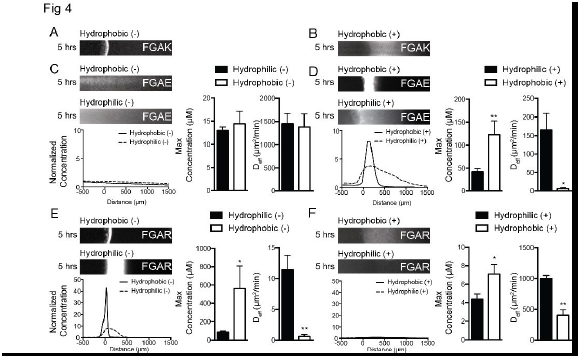
Glutamic acid reverses selectivity of FG-based gels and arginine displays similar selectivity to lysine A)-B) Representative Hydrophobic (−) and Hydrophobic (+) fluorescence images in FGAK gels for comparison to FGAE and FGAR gel selectivity. C) Representative fluorescence images of Hydrophobic (−) and Hydrophilic (−) reporters into FGAE gel with the corresponding concentration profile, max concentration at gel interface, and estimated effective diffusion coefficient. D) Representative fluorescence images of Hydrophobic (+) and Hydrophilic (+) reporters into FGAE gel with the corresponding concentration profile, max concentration at gel interface, and estimated effective diffusion coefficient. E) Representative fluorescence images of Hydrophobic (−) and Hydrophilic (−) reporters into FGAR gel with the corresponding concentration profile, max concentration at gel interface, and estimated effective diffusion coefficient. F) Representative fluorescence images of Hydrophobic (+) and Hydrophilic (+) reporters into FGAR gel with the corresponding concentration profile, max concentration at gel interface, and estimated effective diffusion coefficient. All error bars are standard deviations of at least three independent replicates. * indicates p<0.05. ** indicates p<0.01 between hydrophobic and hydrophilic reporters using unpaired Student’s t-test.

## Sidechain biochemistry as a regulator of assembly

Lastly, we tested how sidechain biochemistry of charged amino acids affect FG sequence function for both self-assembly and molecular recognition of transport molecules. From the Nsp1 consensus sequence analysis, we observed that the predominant cation is lysine rather than arginine, suggesting that the two cations may not be treated equally. Similarly, glutamic acids are conserved closer to FG sequences whereas aspartic acids appear closer to the central region of the spacer sequences and are not as well conserved. To test the role of sidechain differences, we first compare how FGAR gels compared to FGAK. In the diffusion assay, we find that FGAR gels have similar uptake properties when compared to FGAK gels (Figure 4A, E), where the Hydrophobic (−) reporter accumulates and interacts with the gel interface while the Hydrophobic (+) reporter does not accumulate and instead diffuses near freely into the gel (Figure 4E and 4F). These data suggest that that from an electrostatic standpoint, arginines are just as capable as lysines in helping FG sequences differentiate between various types of hydrophobic substrates.

However, mechanically, FGAR forms gels that exhibited an approximate stiffness of 2000 Pa throughout the frequency sweep (Supplementary Figure 4C), which is 4-fold more compliant than FGAK gels. The changes in the mechanical properties of the FGAK and FGAR gels suggest that the lysines and arginines may predominantly affect the structural self-assembly of FG-domains. Indeed, from TEM analysis, we find that FGAR form different structures compared to FGAK (Supplementary Figure 4D). To compare FGAD and FGAE gels, we find that FGAD does not form a gel at the standard 2% (w/v) and forms no repeating structures (Supplementary Figure 4A-B), suggesting that the small chemical changes between glutamic acid and aspartic acid are able to transition the material from a selective gel to a viscous solution. Together, these data show the detailed amino acid biochemistry, in addition to net charge, is an important regulatory parameter in determining the structure of self-assembled FG-domains with minor effects on selectivity.

## Tryptophan interactions are similarly modulated by electrostatics

An essential question in the selectivity of FG-domains is how generalizable the tunability of hydrophobic selectivity is to other natural aromatic amino acids. In previous assays with the designed reporters, we chose phenylalanines as the representative aromatic residue for hydrophobic interactions with FG-domains. In native NTRs such as NTF2 and other transport receptors, tryptophan is commonly used for the transient hydrophobic interactions required for transport. To test if the electrostatic dependence still exists when phenylalanines are converted to tryptophans, we compared two new hydrophobic reporters Hydrophobic (+W) and Hydrophobic (−W) to the original hydrophobic phenylalanine reporters (now termed +F and −F, respectively, for the hydrophobic residue). As seen in Figure 5, the tryptophan reporters are modulated by electrostatic interaction in a similar manner to phenylalanine reporters. We find that the max accumulated fluorescence at the interface and diffusion between Hydrophobic −F and −W reporters are not significantly different (Figure 5A). For cationic reporters, when phenylalanines (+F) are substituted for tryptophans (+W), the reporters exhibit similar max interface properties but tryptophan (+W) reporters show a slight trend toward slower diffusion (Figure 5B), indicating that although both aromatic residues can be modulated by electrostatic interactions, choice of hydrophobic residue will have small but potentially significant differences in transport properties. These results reveal that the phenomenon of electrostatic interactions regulating hydrophobic interactions may be a generalizable phenomenon and is not restricted to phenylalanines. Moreover, the choice of type of non-polar amino acid is now established as another parameter in which selective transport can be tuned.

**Figure 5:**
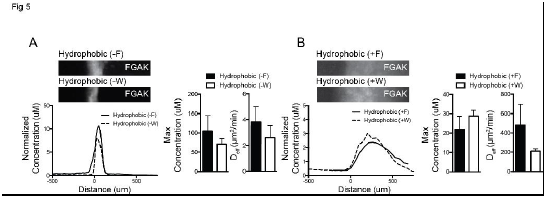
Comparison of selective transport of tryptophan and phenylalanine in FGAK gels A) Comparison of Hydrophobic (−F) and Hydrophobic (−W) reporters into FGAK gels with representative fluorescent images and corresponding concentration profiles. Estimated max interface concentrations and effective diffusion coefficients are presented. B) Comparison of Hydrophobic (+F) and Hydrophobic (+W) reporters into FGAK gels with representative fluorescent images and corresponding concentration profiles. Estimated max interface concentrations and effective diffusion coefficients are presented. All error bars are standard deviations of at least three independent replicates. No significant differences are reported using Student’s t-test.

## Discussion

Our study shows how the environment surrounding FG sequences tunes hydrophobic interactions necessary for correct molecular recognition of hydrophobic substrates. Here, we explicitly show how the presence, placement, and type of charged amino acid are all essential parameters in controlling FG function for these reduced FG repeats. The method of using peptides based on consensus sequences of Nsp1 reveals how complex function can be encoded in minimal sequence space. The work here supports previous theoretical predictions that electrostatic interactions may be an important regulator of nuclear pore selectivity (22) and sequence observations that the bias in the net charge of nuclear transport receptors may be important in NPC transport (21). Moreover, recent reports on how charge influences interactions with hydrophobic surfaces (8,34) emphasize that the interplay between electrostatics and hydrophobicity may be a fundamental process for different biological functions such as molecular recognition and protein folding. Here, we provide an example of how these electrostatics and hydrophobicity can tune the molecular recognition properties of biological supramolecular structures.

Our work shifts the spotlight from analyzing only how hydrophobicity determines nuclear pore selectivity to the underappreciated contribution of electrostatics in molecular recognition (9,11,12,35). The results described here provide insight on how individual hydrophobic FG-sequences may acquire the astounding ability to distinguish a particular subset of hydrophobic substrates using a combination of electrostatic and hydrophobic interactions. Our data also imply that substrate interaction with the FG-containing hydrogel is not a binary effect, but can be tuned by the placement of charge surrounding FG domains to achieve a range of transport behaviors, from complete retention at the interface to free diffusion through the hydrogel, as depicted schematically in Figure 6.

**Figure 6:**
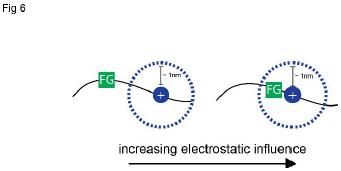
Role of electrostatics in modulating hydrophobic FG function Schematic of the influence of electrostatic on hydrophobic interactions mediated by FG-domains. At physiologically relevant salt concentrations, electrostatic interactions typically have a range of ~1 nm (Debye length), which determines how much a charged residue can influence what hydrophobic substrates FG-domains can recognize and bind to. By moving charged residues further away, the electrostatic effects will be less significant on FG-mediated self-assembly and selective transport.

We recognize that although the peptides can answer questions on how the molecular recognition of FG-sequences can be manipulated by the precise placement of charged amino acids, the short peptide gels cannot recapitulate the unique facilitated diffusion properties of the intact nuclear pore. This likely arises from structural differences, as short peptides form fibers and rods, which are not observed and predicted in the native nuclear pore. Future work to address these challenges include synthesizing longer peptides consisting of multiple consensus repeats to determine how repeat number influences structure and selective recognition. Nevertheless, the current peptides provide unique advantages that are absent in more complex and degenerate systems. For instance, previous studies with intact nucleoporins containing multiple repeats of FG-domains suggest that charge primarily plays a structural role in the cohesion of the self-assembled matrix (36), which results in altered selective properties of the gel. In contrast, peptides with single repeats show that electrostatic interactions directly influence how biological substrates interact with hydrophobic FG sequences for molecular recognition. FG-containing peptides can capture at single amino acid resolution on how the interplay between hydrophobic and electrostatic interactions. These results also support recent work where ImpB was recognized to have varying binding capacities with synthetic FG-containing polyacrylamide gels depending on the charged state of the material (37). We believe this is a useful approach that complements other well-established methods used to understand the selective transport mechanism of NPCs such as the *in vivo* minimal nuclear pore complex (18,19), gel and selectivity analysis of individual nucleoporins (9–11,38), and binding interactions with surface-grafted nucleoporin films (36,39–41). Moreover, we expect that this rational approach using conserved repeating sequences can be expanded to further study other biological gel systems such as mucus, byssal threads, and cartilage, where complex disordered proteins based on repeat units and reversible crosslinking contribute a significant proportion of the material (42–44).

## Materials and Methods

### Sequence analysis

Sequence logo of consensus sequence was generated using Berkeleys’ WebLogo software(45) by aligning 15 repeats according to FSFG as references. Conserved fraction was calculated for positive charges (# cationic residues K or R)/(15 repeats) and similarly for negative charges (# anionic residue E or D)/(15 repeats) at each position indicated.

### NTF2 Expression and Labeling

pQE30-NTF2-6xHis was a gift from the Gorlich lab and transformed into DH5alpha cells for cloning. A cysteine was inserted for fluorescent labeling after the 6x His-tag using standard site directed mutagenesis techniques using the two primers (5’–ctcagctaattaagcttagcagtgatggtgatggtgatgagatctg –3’ and 5’ –cagatctcatcaccatcaccatcactgctaagcttaattagctgag–3’) to form pQE30-NTF2-6xHis-Cys. The W7A-Cys mutant containing the terminal cysteine was created by using pQE30-NTF2-6xHis-Cys and applying standard site direct mutagenesis protocols with the following primers (5’–tgaggagccaatttgttccgcgatcggtttatcacccatg–3’ and 5’–catgggtgataaaccgatcgcggaacaaattggctcctca–3’). Expression of NTF2-Cys and W7A was completed in OverExpress C41(DE3) cells (Lucigen) and purified using standard nickel column purification and ion exchange columns. Labeling of NTF2-Cys and W7A-Cys was completed using Fluorescein-5-maleimide (Thermofischer Scientific, Catalog #F150) using their recommended protocol and gel purified. Labeling efficiency was approximately 50%. Final concentrations of NTF2 and W7A for transport were adjusted to 10 uM concentrations with 10% of the population labeled.

### Peptide and Gel Preparation

Unless specified otherwise, all chemicals were obtained from Sigma Aldrich. Peptides were prepared by MIT’s Koch Institute Biopolymers and Proteomics Facility (Cambridge, MA) and Boston Open Labs LLC (Cambridge, MA). All peptides were HPLC purified unless specified otherwise, desalted using reverse phase HPLC with 0.05% TFA, and lyophilized after synthesis with >95% purity. For fluorescently labeled peptides, a 5-carboxyfluorescein (Anaspec) fluorophore is added to the N-terminus whereas the C-terminus is modified to be an amide. Fluorescent peptides were diluted into 200 mM NaCl with 20 mM HEPES, pH 7 buffer at 10 uM concentration for transport experiments. The gel peptides were all dissolved in 20 mM NaCl, 20 mM HEPES, pH 7 buffer at 2% (w/v). To facilitate solubilization and gel formation, peptides were vortexed for 30 seconds and briefly sonicated in a bath sonicator (Branson 2510) to reduce aggregation.

### Capillary Assay and Analysis

1.5 inch length borosilicate square capillaries with 9 mm cross sectional width (Vitrocom 8290) are loaded by piercing pre-made hydrogels. 10 uM solutions (200 mM NaCl, 20 mM HEPES pH 7) of fluorescent peptides are injected into the capillary and sealed by a 1:1:1 (by weight) mixture of Vaseline, lanolin, and paraffin. Time lapses of peptide diffusion are taken using at 1 minute intervals for up to five hours on a Nikon Ti Eclipse inverted microscope using a Nikon CFI Plan UW 2X or on an AxioObserver D.1 with a EC Plan-Neofluar 1.25x/0.03 WD=3.9 and Hamamatsu C11440-22CU camera. All fluorescence profiles were obtained by averaging the fluorescence intensities across the width of the capillary. Normalized concentration profiles were obtained by normalizing fluorescence intensities to the bath concentration of the capillary at the initial time point. The fluorescence is signal is linear up to 50 uM (Supplementary Figure 5). For concentration profile plotting, the signal is not adjusted past the saturation point and represents the lower bound of the actual concentration of reporters accumulating in the gel. All data presented represents at least three independent replicates. Students t-test was applied to determine p-values between experimental conditions.

Effective diffusion rates were fit by minimizing the squared error of a simulated concentration timecourse in a region of the capillary on the gel side of the interface, over a 100-minute window. To do this, the diffusion equation for the concentration of probe *c*:

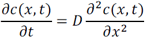

was numerically solved using Matlab’s *pdepe* function (MathWorks; Natick, MA). The initial condition *c*(*x*,0) was set by the concentration profile at the first timepoint used, and the boundary conditions *c*(0,*t*) and *c*(*L*,*t*) (for a fit over length L) were similarly determined by the concentration profiles at the edges of the region of interest. The *D* that minimized the squared difference between simulated and actual concentration profiles was the value reported; minimization of error took place iteratively using a home-written modified gradient descent algorithm. The time period from 150-250 minutes was generally used for fitting, but for particularly fast-diffusing probes (D>1000 μm^2^/min) 50-150 or 30-130 minutes was used. The earlier time period allowed for the fitting to take place before steady-state or pseudo-steady state was reached, which was crucial for fitting to be precise. Code for the diffusion analysis is provided in the supplementary materials.

### Rheological Testing

Experiments were performed on an Anton Paar MCR 302 Rheometer in a cone-plate geometry with a 25mm diameter, 1 degree cone angle, and 51 micron truncation. The temperature was maintained at 25C and evaporation controlled with an H2O-filled solvent trap. To identify the linear regime, amplitude sweeps were conducted at ω = 10 rad/s from γ_0_ = 0.01% to 100% strain. In the linear regime, frequency sweeps were conducted using the previously determined strain amplitude from ω = 100 rad/s to 0.1 rad/s.

### Transmission Electron Microscopy

Images were taken using a JEOL-1200 transmission electron microscope. Gels were formed as aforementioned and spotted onto glow discharged carbon-coated copper-Formvar grids (Ted Pella). Excess liquid and gel were removed using Whatman paper or parafilm. Grids were submerged in 1% uranyl acetate solution, blotted, and air dried for 15 minutes prior to imaging. Images of gels are representative of at least four images taken of each gel type.

### Phenyl-sepharose chromatography

Fluorescent reporters were dissolved in 200 mM NaCl, 20 mM HEPES pH 7 and loaded into high-copy FF 1 mL capacity phenyl-sepharose columns equilibrated with three volumes of 200 mM NaCl, 20 mM HEPES pH 7. For elution, flow rates were set to 1 ml/min with 0.5 mL fraction volumes. The fraction concentrations are representative traces of elution profiles.

### Nile Red Fluorescence Assay

Using the same transport methods, Nile Red dye (Thermo Fischer Scientific, N1142) was loaded into capillaries at 10 ug/ml concentrations in 200 mM NaCl, 20 mM HEPES buffers, pH 7.

## Acknowledgements

This work was supported by the MRSEC Program of the National Science Foundation under award number DMR – 0819762, Defense Threat Reduction Agency under award number HDTRA1-13-1-0038, NSF RO1 R01-EB017755, and NSF Career PHY-1454673. W.G.C. was supported by the NIGMS/NIH Interdepartmental Biotechnology Training Program under T32 GM008334. J.W. was supported in part by the National Science Foundation Graduate Research Fellowship Program under Grant No. 1122374. We wish to thank Prof. Alan Grodzinsky at MIT for his valuable feedback and helpful discussions, and Felice Frankel for the images of the gels.

---

Supplementary Figure 1: Frequency sweep of F→S substitution (FGAK → SGAS) to determine effect of phenylalanines on self-assembly of peptides. The elastic modulus (G’) and loss modulus (G”) are reported, however, the measured values are below the sensitivity of the rheometer using the specified cone-plate geometry.

Supplementary Figure 2: A) Transport of Nile Red dye into FGAK and FGAE gels at the 0 hour and 3 hour timepoint. Fluorescence indicates the dye is able to access hydrophobic environments created by FG-domains within the gels. Images are of representative gels from three independent replicates.

Supplementary Figure 3: Fractionation of hydrophilic reporters compared to their hydrophobic counterparts in phenylsepharose columns. The fluorescence signals of each fraction is collected and normalized to the signal with the highest intensity of emission. For both cationic and anionic reporters, the hydrophobic reporters are eluted later, indicating that they show an increased retention time, and thus stronger binding to phenyl sepharose beads.

Supplementary Figure 4: A) Frequency sweep of FGAD peptide solution with G’ (storage) and G” (loss) moduli reported at 2% (w/v) and corresponding TEM image (B). C) Frequency sweep of FGAR gel with G’ (storage) and G” (loss) moduli reported at 2% (w/v) and corresponding TEM image (D).

Supplementary Figure 5: Quantification of fluorescence signal of fluorophores as a function of concentration. Fluorescence signal is approximately linear up to 50 uM concentrations and saturates by 100 uM. All concentrations are reported according to the experimental curve developed and represent lower values of the actual concentrations if values exceed 100 uM.

## References

1. Stewart M. Molecular mechanism of the nuclear protein import cycle. Nature Reviews Molecular Cell Biology. 2007 Mar 1;8(3):195–208.

2. Stewart M, Baker RP, Bayliss R, Clayton L, Grant RP, Littlewood T, et al. Molecular mechanism of translocation through nuclear pore complexes during nuclear protein import. FEBS Lett. 2001 Jun 8;498(2-3):145–9.

3. Görlich D, Kutay U. Transport between the cell nucleus and the cytoplasm. Annu Rev Cell Dev Biol. Annual Reviews 4139 El Camino Way, P.O. Box 10139, Palo Alto, CA 94303-0139, USA; 1999 Nov;15(1):607–60.

4. Ando D, Zandi R, Kim YW, Colvin M, Rexach M, Gopinathan A. Nuclear pore complex protein sequences determine overall copolymer brush structure and function. Biophys J. 2014 May 6;106(9):1997–2007.

5. Ribbeck K, Görlich D. Kinetic analysis of translocation through nuclear pore complexes. EMBO J. 2001 Mar 15;20(6):1320–30.

6. Ghavami A, Veenhoff LM, van der Giessen E, Onck PR. Probing the Disordered Domain of the Nuclear Pore Complex through Coarse-Grained Molecular Dynamics Simulations. Biophysj. Biophysical Society; 2014 Sep 16;107(6):1393–402.

7. Gamini R, Han W, Stone JE, Schulten K. Assembly of Nsp1 nucleoporins provides insight into nuclear pore complex gating. PLoS Comput Biol. 2014 Mar;10(3):e1003488.

8. Ma CD, Wang C, Acevedo-Vélez C, Gellman SH, Abbott NL. Modulation of hydrophobic interactions by proximally immobilized ions. Nature. 2015 Jan 15;517(7534):347–50.

9. Frey S, Görlich D. A saturated FG-repeat hydrogel can reproduce the permeability properties of nuclear pore complexes. CELL. 2007 Aug 10;130(3):512–23.

10. Frey S, Richter RP, Görlich D. FG-rich repeats of nuclear pore proteins form a three-dimensional meshwork with hydrogel-like properties. Science. 2006 Nov 3;314(5800):815–7.

11. Frey S, Görlich D. FG/FxFG as well as GLFG repeats form a selective permeability barrier with self-healing properties. EMBO J. 2009 Aug 13;28(17):2554–67.

12. Ribbeck K, Görlich D. The permeability barrier of nuclear pore complexes appears to operate via hydrophobic exclusion. EMBO J. 2002 Jun 3;21(11):2664–71.

13. Bayliss R, Littlewood T, Strawn LA, Wente SR, Stewart M. GLFG and FxFG nucleoporins bind to overlapping sites on importin-beta. J Biol Chem. 2002 Dec 27;277(52):50597–606.

14. Bayliss R, Ribbeck K, Akin D, Kent HM, Feldherr CM, Görlich D, et al. Interaction between NTF2 and xFxFG-containing nucleoporins is required to mediate nuclear import of RanGDP. J Mol Biol. 1999 Oct 29;293(3):579–93.

15. Strawn LA, Shen T, Wente SR. The GLFG regions of Nup116p and Nup100p serve as binding sites for both Kap95p and Mex67p at the nuclear pore complex. J Biol Chem. 2001 Mar 2;276(9):6445–52.

16. Grant RP, Neuhaus D, Stewart M. Structural basis for the interaction between the Tap/NXF1 UBA domain and FG nucleoporins at 1A resolution. J Mol Biol. 2003 Feb 21;326(3):849–58.

17. Bednenko J, Cingolani G, Gerace L. Importin beta contains a COOH-terminal nucleoporin binding region important for nuclear transport. J Cell Biol. 2003 Aug 4;162(3):391–401.

18. Strawn LA, Shen T, Shulga N, Goldfarb DS, Wente SR. Minimal nuclear pore complexes define FG repeat domains essential for transport. Nat Cell Biol. 2004 Feb 22;6(3):10–206.

19. Patel SS, Belmont BJ, Sante JM, Rexach MF. Natively unfolded nucleoporins gate protein diffusion across the nuclear pore complex. CELL. 2007 Apr 6;129(1):83–96.

20. Ribbeck K, Kutay U, Paraskeva E, Görlich D. The translocation of transportin–cargo complexes through nuclear pores is independent of both Ran and energy. Current biology. 1999.

21. Colwell LJ, Brenner MP, Ribbeck K. Charge as a selection criterion for translocation through the nuclear pore complex. PLoS Comput Biol. 2010 Apr;6(4):e1000747.

22. Tagliazucchi M, Peleg O, Kröger M, Rabin Y, Szleifer I. Effect of charge, hydrophobicity, and sequence of nucleoporins on the translocation of model particles through the nuclear pore complex. Proceedings of the National Academy of Sciences. 2013 Feb 26;110(9):3363–8.

23. Ando D, Colvin M, Rexach M, Gopinathan A. Physical Motif Clustering within Intrinsically Disordered Nucleoporin Sequences Reveals Universal Functional Features. PLoS ONE. 2013;8(9):e73831.

24. Yamada J, Phillips JL, Patel S, Goldfien G, Calestagne-Morelli A, Huang H, et al. A bimodal distribution of two distinct categories of intrinsically disordered structures with separate functions in FG nucleoporins. Mol Cell Proteomics. 2010 Oct;9(10):2205–24.

25. Zhang S. Fabrication of novel biomaterials through molecular self-assembly. Nature Biotechnology. 2003 Oct;21(10):1171–8.

26. Hinman MB, Jones JA, Lewis RV. Synthetic spider silk: a modular fiber. Trends Biotechnol. 2000 Sep;18(9):374–9.

27. Xu M, Lewis RV. Structure of a protein superfiber: spider dragline silk. Proceedings of the National Academy of Sciences. 1990 Sep 15;87(18):7120–4.

28. Meyer DE, Chilkoti A. Quantification of the effects of chain length and concentration on the thermal behavior of elastin-like polypeptides. Biomacromolecules. 2004 May;5(3):846–51.

29. Nettles DL, Chilkoti A, Setton LA. Applications of elastin-like polypeptides in tissue engineering. Adv Drug Deliv Rev. 2010 Dec;62(15):1479–85.

30. Wright ER, Conticello VP. Self-assembly of block copolymers derived from elastin-mimetic polypeptide sequences. Adv Drug Deliv Rev. 2002 Oct 18;54(8):1057–73.

31. Hurt EC. A novel nucleoskeletal-like protein located at the nuclear periphery is required for the life cycle of Saccharomyces cerevisiae. EMBO J. 1988 Dec 20;7(13):4323–34.

32. Denning DP. Disorder in the nuclear pore complex: The FG repeat regions of nucleoporins are natively unfolded. Proceedings of the National Academy of Sciences. 2003 Feb 25;100(5):2450–5.

33. Denning DP, Uversky V, Patel SS, Fink AL. The Saccharomyces cerevisiae nucleoporin Nup2p is a natively unfolded protein. … of Biological Chemistry. 2002.

34. Chen S, Itoh Y, Masuda T, Shimizu S, Zhao J, Ma J, et al. Ionic interactions. Subnanoscale hydrophobic modulation of salt bridges in aqueous media. Science. 2015 May 1;348(6234):555–9.

35. Hülsmann BB, Labokha AA, Görlich D. The permeability of reconstituted nuclear pores provides direct evidence for the selective phase model. CELL. 2012 Aug 17;150(4):738–51.

36. Ader C, Frey S, Maas W, Schmidt HB, Görlich D, Baldus M. Amyloid-like interactions within nucleoporin FG hydrogels. Proceedings of the National Academy of Sciences. 2010 Apr 6;107(14):6281–5.

37. Friedman AK, Baker LA. Synthetic hydrogel mimics of the nuclear pore complex display selectivity dependent on FG-repeat concentration and electrostatics. - PubMed - NCBI. Soft Matter. 2016.

38. Labokha AA, Gradmann S, Frey S, Hülsmann BB, Urlaub H, Baldus M, et al. Systematic analysis of barrier-forming FG hydrogels from Xenopus nuclear pore complexes. EMBO J. 2013 Jan 23;32(2):204–18.

39. Eisele NB, Labokha AA, Frey S, Görlich D, Richter RP. Cohesiveness tunes assembly and morphology of FG nucleoporin domain meshworks - Implications for nuclear pore permeability. Biophys J. 2013 Oct 15;105(8):1860–70.

40. Eisele NB, Frey S, Piehler J, Görlich D, Richter RP. Ultrathin nucleoporin phenylalanine-glycine repeat films and their interaction with nuclear transport receptors. EMBO Rep. 2010 May;11(5):366–72.

41. Zahn R, Osmanović D, Ehret S, Callis CA, Frey S, Stewart M, et al. A physical model describing the interaction of nuclear transport receptors with FG nucleoporin domain assemblies. Elife. eLife Sciences Publications Limited; 2016 Apr 8;5:e14119.

42. Holten-Andersen N, Zhao H, Waite JH. Stiff coatings on compliant biofibers: the cuticle of Mytilus californianus byssal threads. Biochemistry. 2009 Mar 31;48(12):2752–9.

43. Waller KA, Zhang LX, Elsaid KA, Fleming BC, Warman ML, Jay GD. Role of lubricin and boundary lubrication in the prevention of chondrocyte apoptosis. Proc Natl Acad Sci USA. 2013 Mar 25;:-.

44. Rose MC, Voynow JA. Respiratory tract mucin genes and mucin glycoproteins in health and disease. Physiol Rev. 2006 Jan;86(1):245–78.

45. Crooks GE, Hon G, Chandonia J-M, Brenner SE. WebLogo: a sequence logo generator. Genome Res. 2004 Jun;14(6):1188–90.

